# Estrogen Receptor Beta Localized on Ventral Tegmental Area Dopamine Neurons Regulates Nicotine Self-Administration Acquisition in Ovary-Intact Female Rats

**DOI:** 10.1101/2025.07.30.667767

**Authors:** Ashley M. White, Kathleen R. McNealy, Shailesh N. Khatri, Sally Pauss, Brandon Henderson, Heather A. Bimonte-Nelson, Terry D. Hinds, Cassandra D. Gipson

**Author notes:** **Corresponding Author:** Cassandra D. Gipson, Associate Professor, Department of Pharmacology and Nutritional Sciences, University of Kentucky, 535 CT Wethington Building, 900 S. Limestone, Lexington, KY 40536.

## Abstract

Women exhibit greater nicotine use vulnerability than men. High estradiol (E2) exacerbates nicotine use outcomes in women, effects which have been modeled in preclinical nicotine self-administration (SA) studies. Nicotine SA is maintained by dopamine (DA) release from the ventral tegmental area (VTA) to the nucleus accumbens (NA). E2 exerts its effects by binding to estrogen receptors (ER), including ERα, ERβ, and G-protein coupled ER-1 (GPER-1)s. E2 action at ERs specifically has been shown to potentiate DA neuronal excitability within the VTA. Further, we have shown that ovariectomy decreases both nicotine use during SA and VTA ERβ protein. Despite clear evidence of mechanistic relationships between E2, ERs, DA, and nicotine, no studies to date have functionally evaluated the specific role of ERβ located on VTA DA cells in driving nicotine consumption during SA in females. There are currently no tools that allow for evaluations of relationships between nicotine neurobiology and ERs with cell-type specificity, as ERβ is also localized on other (non-DA) cell types within the VTA. As such, the goals of the present studies were (1) to build and validate a novel adeno-associated viral construct that produces long-term knockdown of ERβ specifically on VTA DA neurons, and (2) to determine if VTA DA ERβ viral knockdown reduces nicotine SA in ovary-intact female rats. Here we show that ERβ regulates VTA DA neuron excitability, and that ERβ knockdown in VTA DA neurons reduces DA neuron firing frequency. We also show that VTA ERβ knockdown in DA neurons reduces nicotine SA acquisition in ovary-intact female rats. Together, our results demonstrate a critical role of ERβ in driving nicotine use in females, underscoring the need for future studies to evaluate neurobehavioral mechanisms of smoking through the lens of sex differences.

## Introduction

Nicotine, the primary active alkaloid in tobacco, is the primary addictive agent responsible for the abuse liability of cigarettes and other tobacco products (e.g., e-cigarettes Stolerman and Jarvis [1]). Use of nicotine-containing products remains the leading cause of preventable death and disease, with cigarettes alone contributing to over 480,000 deaths per year in the United States [2]. Smoking rates are decreasing slower for women than men [3]. Women experience greater nicotine use vulnerability than men, insofar that they become addicted to nicotine faster than men and exhibit higher rates of smoking relapse [4; 5]. In turn, long-term smoking cessation is more difficult to achieve in women than in men [6; 7; 8; 9; 10], likely resulting from commonly used smoking cessation pharmacotherapies (e.g., nicotine patch and varenicline [11; 12] being less effective for women. Additionally, men who smoke demonstrate a choice preference for lower nicotine doses in an intravenous nicotine self-administration (SA) task while women who smoke do not display a preference for lower doses in this task [13].

There is clear clinical evidence for sex hormones estradiol and progesterone playing a role in enhanced nicotine use vulnerability amongst women. For example, cessation attempts are less successful in the follicular phase of the menstrual when estradiol is high, and more successful in the luteal phase when progesterone is high [14; 15; 16]. Hormonal fluctuations across the menstrual cycle also contribute to similar alterations in *ad libitum* smoking [17; 18; 19], economic demand for cigarettes [20], subjective effects of nicotine [21; 22], and sensitivity to smoking cues [23]. This and other converging clinical evidence across numerous nicotine use outcomes supports enhanced nicotine use vulnerability by 17β-estradiol (E2) and protective effects of progesterone. These consistent sex and hormone-related differences at the clinical level warrant preclinical evaluations of the female-specific neurobehavioral mechanisms that may contribute to women’s unique experiences with nicotine.

Preclinical studies have uncovered a wealth of information regarding specific neurobiological circuits [14; 15] and cellular mechanisms [16; 24] underlying nicotine use. Through this seminal work, we know that nicotine engenders self-administration (SA) behavior through modulating dopamine (DA) release from the ventral tegmental area (VTA) to the nucleus accumbens (NA) as a result of nicotine agonism at β2-containing nicotinic acetylcholine receptors (nAChRs) [16]. The VTA is a heterogeneous brain region composed of 60-65% DA neurons (with 30-35% being either local GABAergic interneurons, projection neurons [25], or local glutamatergic neurons that reside within the medial VTA [26]). VTA DA neurons project to the NA and modulate the function of GABAergic medium spiny neurons (MSNs) located within the terminal region (these cells were found were in a metaplastic state following OVX and nicotine SA in our prior study [27]) through DA binding to D1- or D2-receptors [28] . Akin to other addictive drugs, nicotine increases DA within the NA [29]. DA from VTA projecting DAergic neurons is critical in driving nicotine SA [30], evidenced by DA antagonism [31; 32], and lesions [33] of the VTA attenuating nicotine SA.

Notably, a large majority of the seminal studies contributing to our knowledge of what drives nicotine use acquisition have been limited to evaluations in male non-human animal models [30; 31; 32; 33; 34] As with the clinical literature, there is substantial evidence that nicotine use outcomes vary by sex and hormonal state in preclinical models. Females generally self-administer more nicotine than males [35; 36; 37]. As detailed in the following paragraph, we and others have also found evidence of sex steroid modulation of nicotine SA and neurobiology [38; 39; 40; 41]. Thus, there is a large gap in our understanding of female-specific neurobiological mechanisms altering acquisition of nicotine intake more generally, or the mechanisms by which neuroactive steroid hormones impact the brain reward pathway and nicotine use in females more specifically. Thus, preclinical studies are needed that can reveal the relationships between hormonal impacts on neurobiology and nicotine consumption behavior in females.

Removal of the ovaries via ovariectomy (OVX) results in a precipitous loss of ovary-derived steroid hormones and persistent diestrus (acyclicity [42]). Studies with OVX rats thus provide evidence for the role of acute sex steroid hormone modulation of nicotine use outcomes preclinically. We showed that in female rats, OVX decreases nicotine consumption in a SA task but potentiates NA glutamate plasticity [34]. To identify specific hormone influences, we further examined the effects of exogenous E2 supplementation. E2 replacement partially reversed OVX-induced decreases in nicotine SA and potentiation of NA glutamate plasticity in a nicotine-specific fashion [27]. We have further shown that OVX results in reductions in estrogen receptor beta (ERβ) protein expression within the VTA following nicotine SA [43]. Thus, VTA ERβ may be an important target by which ovary-derived steroid hormones act to drive nicotine use in females. Further, it is biologically feasible that E2 can impact nicotine use vulnerability through interactions with ERs within the VTA-->NA DA pathway, given that E2 directly regulates gene expression and cell signaling within the striatum [44; 45; 46]. When animals are in diestrus (when E2 levels are rising), there is increased sensitivity of VTA DA neurons [47]. This result supports that during estrous cycle phases when E2 levels are rising, DA neurons in the VTA more readily fire, which may then increase DA in the NA. E2 exerts its effects through binding to ERs [48] and subsequently increasing dopamine receptor density, enhancing firing of dopaminergic neurons, increasing dopamine transmission, and enhancing dopamine release [48; 49]. More specifically, E2 action at ERs has been shown to potentiate DA neuronal excitability within the VTA [47].

Despite these well-established mechanistic relationships between E2, ERs, DA, and nicotine, no studies to date have functionally evaluated the specific role of ERβ located on VTA DA cells in driving acquisition of nicotine SA in females. While our prior studies utilizing OVX demonstrated significant reductions in nicotine SA that are attenuated by chronic E2 and ethinyl estradiol (EE; a synthetic estrogen) treatment [27; 43], this procedure does not allow for evaluations of the role of specific ERs in modulation of nicotine SA. We [27] and others [50; 51] have shown that OVX itself modulates protein expression and/or mRNA of all ERs in brain tissue. The direction of this modulation occurs differentially based on ER subtype (including ERα [52], ERβ [50], and GPER1) and brain region. Additionally, E2 and EE both bind to ERα and ERβ. E2 binds with roughly equal affinity to ERα and ERβ, and EE has roughly twice the affinity for ERα and half the affinity for ERβ than does E2 [53]. Thus, prior work with exogenous estrogen treatment in OVX rats has provided crucial preliminary evidence for modulation of nicotine SA and neurobiology by neuroactive steroid hormones and estrogens/ERs. However, this evidence is limited in its translation to ovary-intact/freely cycling organisms (e.g., premenopausal women) and in its mechanistic specificity to ERβ.

In addition to limitations from prior work with OVX rats, there are currently no tools that allow for evaluations of relationships between nicotine neurobiology and ERs with cell-type specificity, as ERβ is also localized on other (non-DA) cell types within the VTA. For example, systemic treatment with ERβ antagonists such as 4-[2-Phenyl-5,7-bis(trifluoromethyl)pyrazolo[1,5-a]pyrimidin-3-yl]phenol (PHTPP) can elucidate effects of ERβ signaling on behavioral and neurobiological outcomes. However, in addition to the central nervous system, ERβs are located throughout the body [54], and systemic PHTPP targets ERβ both inside and outside of the central nervous system [55]. Intra-VTA injection of ERβ agonists (such as diarylpropionitrile [DPN]) and antagonists (such as PHTPP or methyl-piperidino-pyrazole [MPP]) can begin to address this limitation. However, as mentioned earlier, ERβ is also localized on other (non-DA) cell types within the VTA. Thus, while intra-VTA pharmacological modulation of ERβ can explicate the role of VTA ERβ signaling, it cannot isolate effects of ERβ specifically on DA neurons. Further, this approach requires repeated intracranial injections which can result in tissue trauma that produces extraneous effects and limits study duration [56]. Thus, for a rigorous and mechanistically specific assessment of the role of ERβ modulation of DA signaling on nicotine SA, we need an approach that specifically and constitutively modulates ERβ receptors on DA neurons within the VTA. No such tools that meet these criteria are currently available. Given this gap and the scarcity of research examining mechanisms of sex steroid hormone alterations of nicotine SA in females, the goals of present studies were two-fold: (1) to build and validate a novel adeno-associated virus (AAV) that encapsulates a short hairpin RNA (shRNA) to produce long-term knockdown of ERβ specifically on VTA DA neurons, and (2) to determine if this VTA DA ERβ viral knockdown reduces nicotine SA in ovary-intact female rats. We hypothesized that akin to our prior effects with OVX, viral knockdown of ERβ DA cells in the VTA would suppress nicotine SA in females.

## Materials and Methods

### 2.1 Subjects

For the electrophysiology ERβ antagonism assay, 3 ovary-intact nicotine-naive female C57BL/6J mice (3 months old) were used (Marshall University). For the *in vivo* viral validation and behavioral studies, 6 ovary-intact female rats (3 per group) were used for patch clamp electrophysiology recordings (see Section 2.5) and 16 ovary intact female Long-Evans rats (Charles River; 200-250g with an arrival of approximately 8 weeks) were used for SA (N=8 per group; see section 2.8; University of Kentucky). Rats were individually housed on a 12:12 reverse light cycle with *ad libitum* access to food and water prior to experimental procedures. Animals were handled daily upon arrival. XX rats were excluded from all analyses due to catheter patency failure. All rat use practices were approved by the Institutional Animal Care and Use Committee of University of Kentucky (UK; Protocol # 2020-3438) and Marshall University (Protocol #664).

### 2.2 Drugs

Nicotine tartrate salt (MP Biomedicals, LLC, Solon, OH, USA) was dissolved in 0.9% saline, and pH was adjusted to 7.4. All nicotine doses are reported in base form. Ketamine (Akorn Animal Health, Lake Forest, IL), xylazine (Akorn Animal Health, Lake Forest, IL), cefazolin (Qilu Pharmaceutical, Shandong, China), meloxicam (Norbrook Inc., Overbrook, KS), and heparin (Sagent Pharmaceuticals, Schaumburg, IL), PHTPP (Sigma-Aldrich, St. Louis, MO), and DPN (Tocris Bioscience, Bristol, UK) were applied/administered as reported below.

### 2.3 Patch Clamp Electrophysiology with ERβ Antagonism in Mice

For the data presented in **Figures 1A-F**, mice were exposed to CO2 and then a cardiac perfusion was performed using ice-cold NMDG-based artificial cerebrospinal fluid (NMDG-ACSF) saturated with 95%/5% O2/CO2 (carbogen) containing (in mM): 93 NMDG, 2.5 KCl, 1.2 NaH2PO4, 10 MgSO4, 0.4 CaCl2, 30 NaHCO3, 5 Na-ascorbate, 3 Na-pyruvate, 2 thiourea, and 25 glucose. Brains were placed in agarose for slicing with a Compresstome® VF-300-OZ (Precisionary Instruments). Coronal brain sections (300 µm) were cut into cold carbogenated NMDG-ACSF to obtain slices containing the VTA (target bregma -2.8 to -3.4 mm). Slices recovered at 32°C in carbogenated NMDG-ACSF for 12-15 min. Next, slices were transferred to standard ACSF containing (mM): 125 NaCl, 2.5 KCl, 1.2 NaH2PO4, 1.2 MgCl2, 2.4 CaCl2, 26 NaHCO3, and 11 glucose for one hour at 32°C. One hour later, slices were transferred to the recording chamber and perfused with carbogenated ACSF (1.5 - 2.0 ml/min) at room temperature.

**Figure 1.**
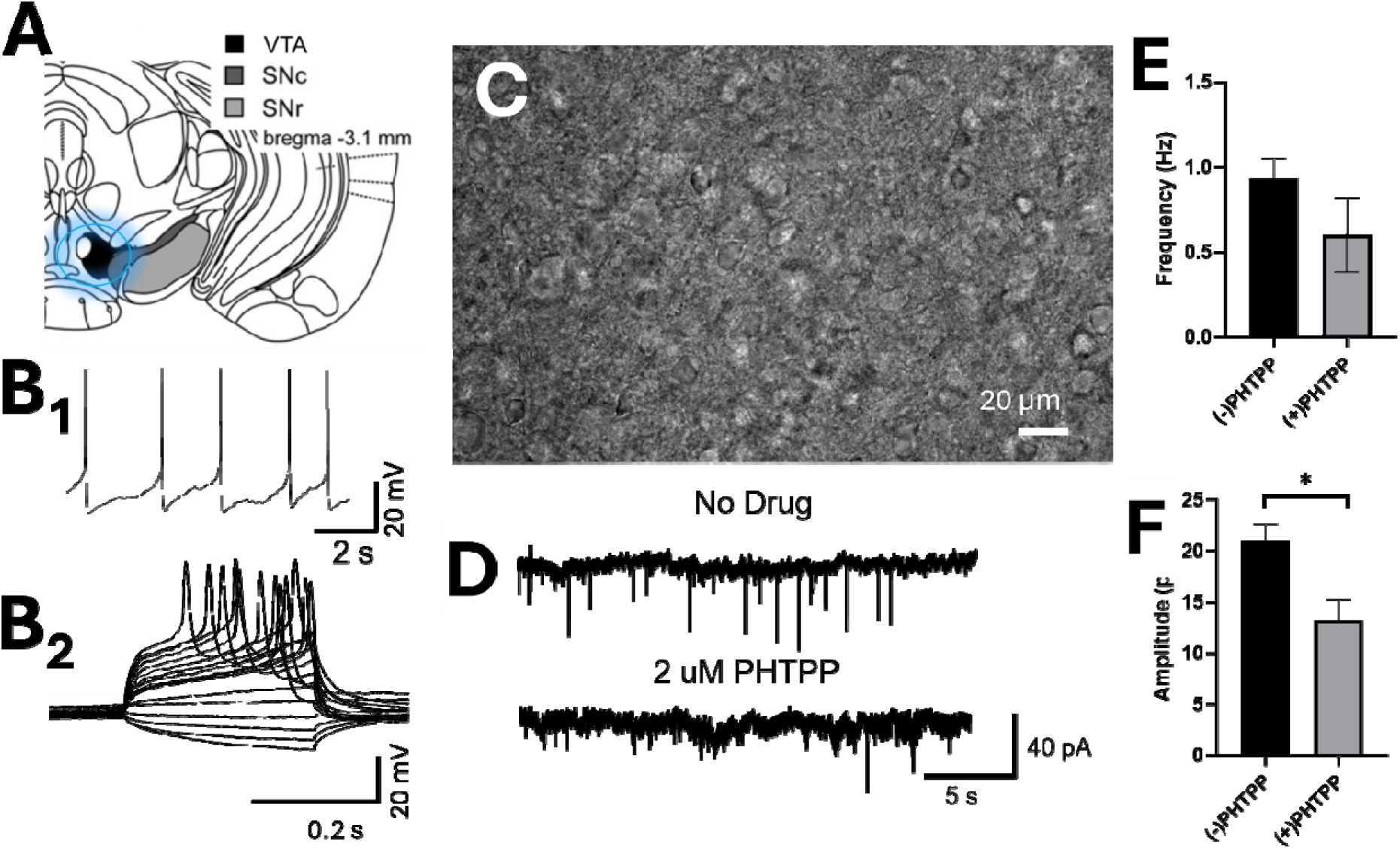
ERβ Antagonism Reduces DA Firing Frequency in the VTA. **(A)** Electrophysiological targeting of the VTA. **(B-_1_-)** Sample recordings of baseline firing and current injections to measure rheobase **(B_2_)** of mouse VTA DA neurons. **(C)** DA cell bodies identified for patch clamp. **(D)** Traces from DA neurons with either no drug or PHTPP (2 µM). **(E)** PHTPP reduced frequency and **(F)** amplitude of sEPSCs in VTA DA neurons. Mean±SEM, \**p*<0.05, one-tailed *t* test. Red arrow=cell soma. All cells were positive for hyperpolarizing *I_h_*, and all had firing frequencies within typical DA firing.

Neurons were visualized with an Axio Examiner A1 (Zeiss) equipped with an Axiocam 702 mono (**Figure 1C**). We recorded electrophysiological signals with an Integrated Patch-Clamp Amplifier (Sutter) using previously described methods. Electrodes had resistances of 4 – 10 MΩ when filled with intrapipette solution (in mM): 135 K gluconate, 5 KCl, 5 EGTA, 0.5 CaCl2, 10 HEPES, 2 Mg-ATP, and 0.1 GTP. Recordings were sampled at ≥10 KHz. The junction potential between patch pipette and bath solutions was nulled just before gigaseal formation. Series resistance was monitored without compensation throughout experiments using SutterPatch software and recordings were terminated if series resistance changed by >20%. In whole-cell recordings, recordings were made after 5 minutes to provide sufficient time for interchange of intrapipette solution with intracellular components.

Spontaneous excitatory post-synaptic currents (EPSCs) were recorded with a holding potential of -70 mV. PHTPP was applied to brain slices through perfusion at a concentration of 2 μM. Prior to drug application, neurons were recorded for 2 – 5 minutes. PHTPP was perfused onto the slices for 2 minutes and then sEPSCs were recorded for 2 – 5 minutes.

### 2.4 Surgical Procedures

One week after acclimation to our animal facility, rats underwent jugular vein catheterization (JVC) surgery then stereotactic craniotomies for viral infusions. Animals were first anesthetized with intramuscular (i.m.) ketamine (80-100 mg/kg) and xylazine (8 mg/kg) and implanted with an indwelling jugular vein catheter as described previously [57]. We made a small incision on the rat’s neck to visualize the jugular vein, followed but one small incision between the scapulae. A 13 cm catheter with a silicon anchor (Instech, Plymouth Meeting, PA) was tunneled subcutaneously such that the anchored side was inserted into the jugular vein and the other end was attached to a backport for SA (Instech, Plymouth Meeting, PA). Immediately after catheter implantation, rats were maintained on isofluorane anesthesia. We then performed stereotactic craniotomies. We made a small incision to visualize the rat’s skull. After identifying bregma, the VTA was located using the following coordinates: A/P: -5.0; M/L: 0.6; D/V: -8.5, where two drill holes were placed at the top of the cranium. Bilateral infusions of the ERβ-shRNA AAV5 or scramble control AAV5 were delivered into the VTA with a Hamilton syringe at a rate of 0.1 µl/min, ∼10^12^ vg/ml. Rats were given a 7-day recovery period prior to experimental procedures. On the day of surgery and for the first two days of surgical recovery, rats were administered meloxicam (1 mg/kg subcutaneously [s.c.]) for post-operative pain management. Additionally, we administered antibiotic cefazolin (100 mg/kg, intravenous [i.v.]) immediately after surgery and throughout the seven-day recovery period to prevent post-surgical infection. The 7-day recovery period allows for adequate post-surgical healing and stable viral infection before beginning experimental procedures. We maintained catheter patency throughout SA by daily pre- and post-SA session administration of 0.1 mL of heparinized saline (100 USP units/mL, i.v.).

### 2.5 Patch Clamp Electrophysiology for Validation of ERβ Knockdown

#### Slice preparation

Rats expressing appropriate AAV (**Figure 2A and B**) were anesthetized with isoflurane and rapidly decapitated, akin to other studies from our lab and others [57; 58; 59]. Brains were quickly removed and placed into ice-cold cutting solution (cutting aCSF) containing (in mM): 92 NMDG, 2.5 KCl, 1.25 NaH2PO4, 30 NaHCO3, 20 HEPES, 25 glucose, 2 thiourea, 5 Na-ascorbate, 3 Na-pyruvate, 0.5 CaCl2·2H2O, and 10 MgSO4·7H2O and saturated with carbogen (95% O2 / 5% CO2), pH:7.3-7.4, and osmolarity: 295 ± 5 mOsm. Coronal slices of 300 μm thickness containing the VTA were made on a vibratome (Leica, VT1000S), through the slicing process the brain was submerged in the same cutting solution that was constantly oxygenated. Slices were immediately transferred to an oxygenated holding chamber containing aCSF maintained at 32-34C for ∼30 mins. While in recovery, NaCl was gradually added to the holding chamber as described in [60]. Post recovery, slices were transferred to another holding chamber maintained at room temperature containing holding solution (in mM: 92 NaCl, 2.5 KCl, 1.25 NaH2PO4, 30 NaHCO3, 20 HEPES, 25 glucose, 2 thiourea, 5 Na-ascorbate, 3Na-pyruvate, 2 CaCl2·2H2O, and 2 MgSO4·7H2O, 295 ± 5 mOsm and pH 7.3-7.40). Slices were maintained at room temperature for 4-6 h and used for recording.

**Figure 2.**
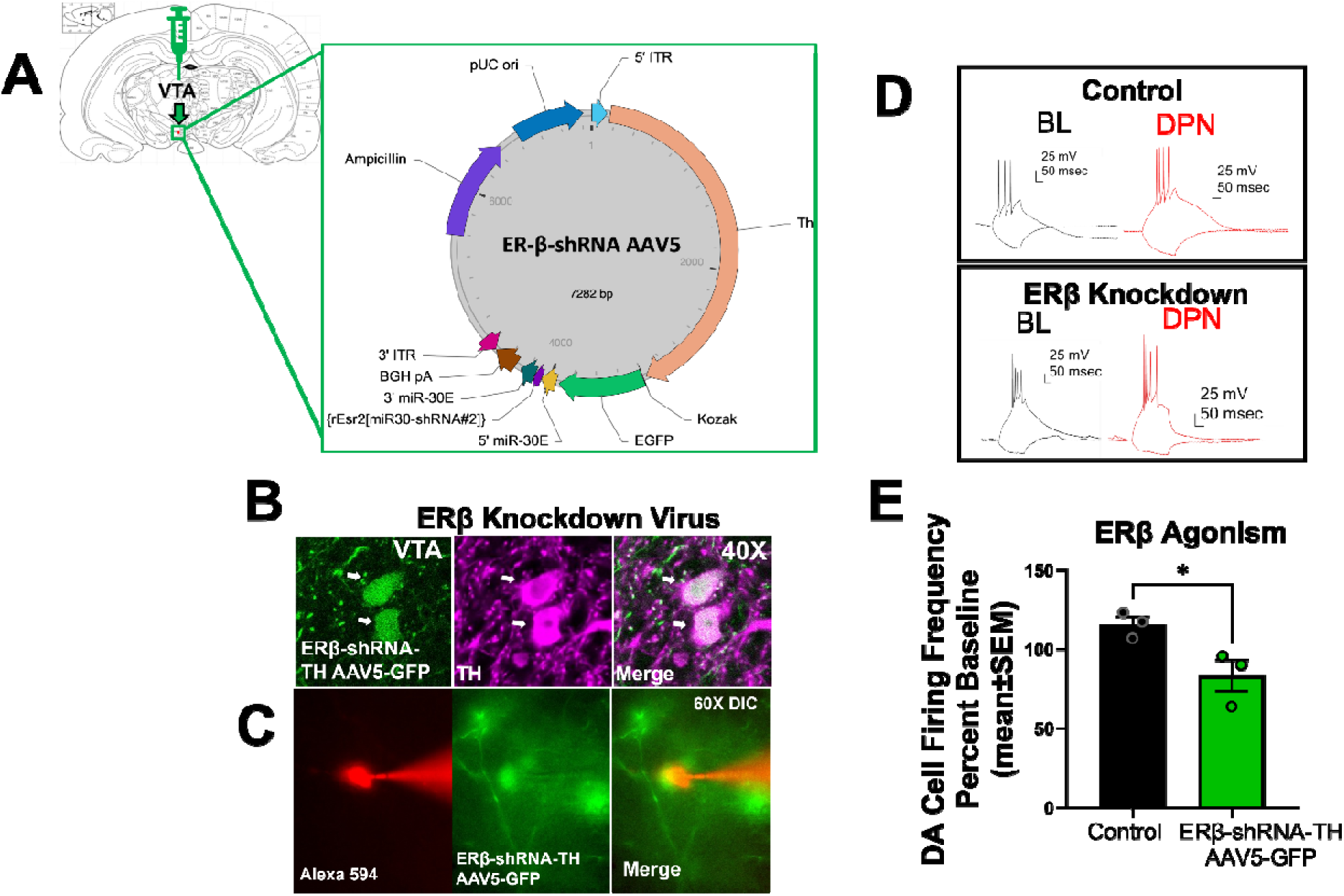
VTA DA ERβ Knockdown Reduces DA Firing Frequency. **(A)** Map of ERβ-shRNA AAV5 microinjected into the VTA. The AAV5 construct contains a tyrosine hydroxylase (TH) promoter that drives green fluorescent protein (EGFP) and the Esr2-shRNA to suppress rat ERβ in VTA DA cells. **(B)** GFP from the ERβ knockdown virus co-localizes with TH in the VTA. **(C)** Patching virus-infected DA neurons in the VTA. **(D)** Traces from ERβ knockdown and scramble control-infected DA neurons. **(E)** ERβ knockdown significantly reduces % baseline (no drug) DA cell firing frequency when DPN is bath applied compared to a no virus control condition. Unpaired *t* test; \**p*<0.05. BL=baseline in D.

#### Recording

For recording, slices were gently transferred to a recording chamber that was continuously perfused with aCSF at flow rate of 1-2 ml/min. The VTA was visually located using a 10X air objective on an Olympus BX51WI fixed stage upright microscope and confirmed by expression of GFP (**Figure 2C**). The Th-positive, GFP-expressing cells were identified by switching to 60X water immersion objective and briefly illuminating slices with a blue LED (approx. 470-480 nm) via an upright microscope equipped with a GFP filter cube, allowing for the direct visualization of fluorescent TH-GFP neurons. Once a target cell was identified, the LED illumination was switched off, and the microscope transitioned to infrared differential interference contrast (IR-DIC) equipped with ORCA-spark digital CMOS camera (Hamamatzu, NJ, USA), for precise pipette positioning and subsequent patch-clamp recording.

Whole-cell recordings were made from the soma of TH^+^ GFP-expressing cells (**Figure 2C**) using recording pipettes (4-7 mΩ) after establishing a giga-ohm seal (resistance range: 1-10 GΩ). Recording pipettes were made pulling borosilicate glass pipettes (Sutter instruments, CA, USA) on Narishige pipette puller (Model PC-100, Automate scientific, CA, USA) and filled with an intracellular solution containing (in mmol/L): K-gluconate, 135; NaCl, 12; K-EGTA, 1; HEPES, 10; Mg-ATP, 2 and tris-GTP, 0.38. Osmolarity was adjusted to 285±5 mOsm, and pH was adjusted to 7.30 ± 0.01. Additionally, Alexa Fluor 594 (20 µM) was added to the internal solution for post-patch analysis if necessary. Upon membrane rupture, the cell membrane potential was held at -70mV. For current-clamp recordings, Action potentials were evoked by depolarizing current injections (200 ms, 700 pA) through the patch pipette from a hyperpolarized baseline membrane potential (-70 mV). Electrophysiological currents were recorded with a Multiclamp 700B amplifier (Molecular Devices, CA), filtered at 10 kHz and digitized at 50LJkHz. Data were collected using pCLAMP software (Molecular Devices).

Recordings without a stable holding current prior to drug application were discarded. Only cells that exhibited overshooting, thin action potentials, normal resting membrane potentials, and changes in uncompensated access resistance less than 20mΩ were included in analyses. Access resistance, membrane resistance, and resting membrane potential (monitored in zero current mode) were continuously monitored throughout recordings to ensure cell viability. The recording software pClamp 10 (molecular devices) was used for conducting recordings. All recordings were digitized at 10 kHz and saved using the digidata interface (Axon Instruments), and analyzed offline using IgorPro (Wavemetrics, Lake Oswego, OR) with the Neuromatic toolkit [61].

### 2.6 Operant Conditioning Chambers

Experimental sessions occurred in operant chambers (MED Associates, St. Albans, VT) within sound-attenuating cubicles. Two levers, one active and one inactive, were extended at the beginning of each session. Above each lever was a stimulus light and each chamber contained a house light. Infusion pumps (MED Associates) were located outside of each chamber. Ventilation fans were provided in each sound-attenuating chamber. Experimental events were programmed and recorded by MED-PC software (MED Associates) on a computer in the experimental room. All operant chambers have been previously described in detail [62; 63].

### 2.7 Food Training Procedures

Following surgical recovery, rats were food restricted to 85% of their free fed body weight. They completed a fifteen-hour food training procedure on a fixed-ratio 1 (FR1) schedule of reinforcement where one active lever press resulted in delivered of a 45 mg food pellet in the operant chamber. There was no programmed consequence of pressing the inactive lever. Animals were required to meet a 2:1 active to inactive lever pressing ratio with a minimum of 200 active lever presses to move onto SA procedures.

### 2.8 SA Procedures

An experimental timeline can be seen in **Figure 3A**. Rats remained food restricted to 85% of their free-fed body weight during self-administration. Infusions of either 0.06 mg/kg/infusion nicotine or saline (0.1 mL/infusion) were delivered across 5.9 s following one response on the active lever (FR-1). Upon an active lever press, lights above both levers were illuminated, and a tone (2900 Hz) was presented simultaneously with drug infusion. Upon completion of the infusion, the tone and lights ceased. Infusions were followed by a 20-s timeout period, during which active lever responses were recorded but produced no consequences. An inactive lever was present at all times but produced no consequences when pressed. All sessions were 2 hours in duration. Rats completed a minimum of 11 sessions with a minimum of 10 active lever presses at a 2:1 active to inactive ratio. Criterion for acquisition was such that rats had to meet this 2:1 active to inactive ratio for three consecutive self-administration sessions.

**Figure 3.**
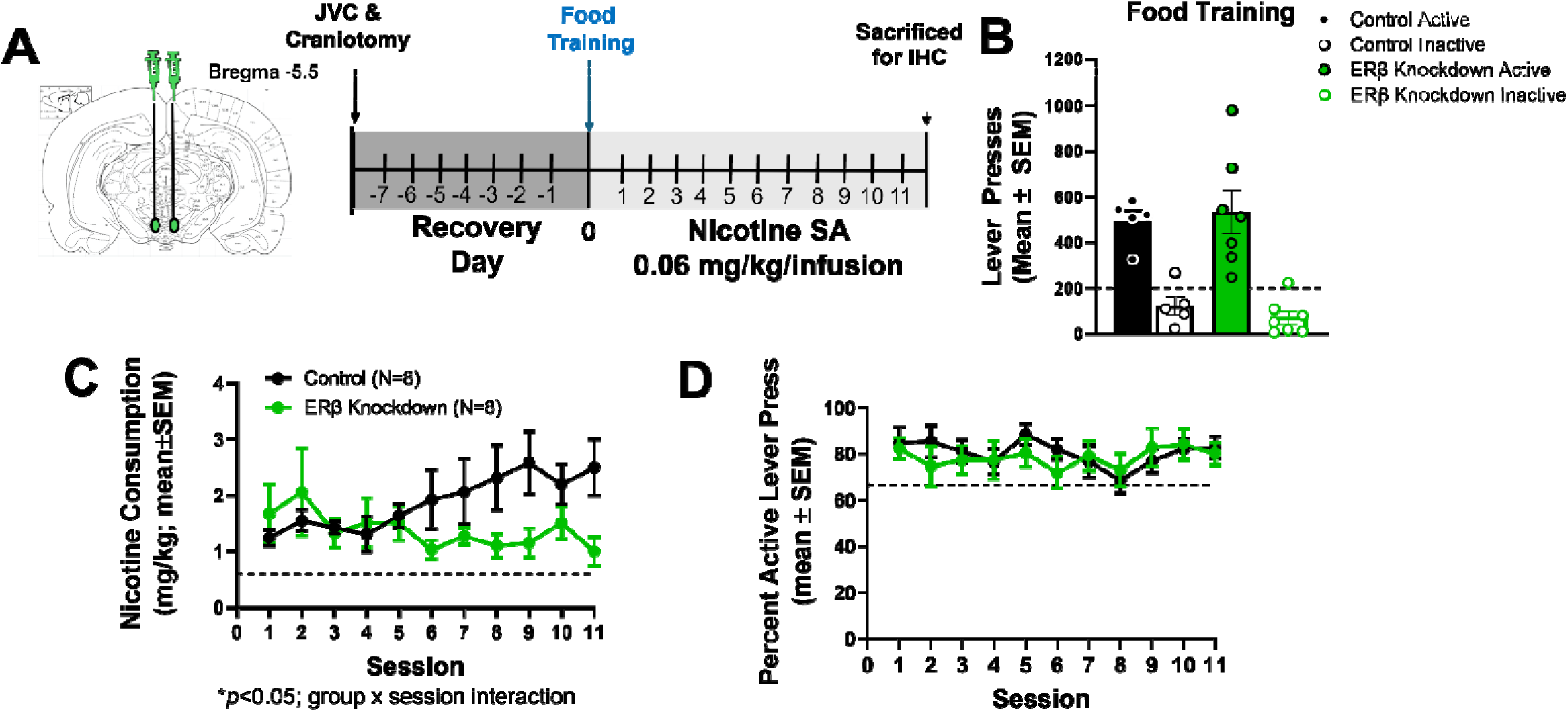
VTA DA ERβ Knockdown Reduces Nicotine Consumption. **(A)** Depiction of virus injection and timeline of surgery and nicotine SA procedures. **(B)** There were no differences in preliminary food training between virus control groups prior to nicotine SA (dotted line = 200 active lever press minimum criterion). **(C)** ERβ knockdown suppressed nicotine consumption in later SA sessions. Dotted line = acquisition criterion set at 0.6 mg/kg or 10 infusions. **(D)** Lever discrimination (active/active+inactive) during SA is not impacted by ERβ knockdown. JVC=jugular vein catheter. *\*p*<0.05 in E.

### 2.9 Data analysis

#### Electrophysiology

For patch clamp electrophysiology in mice (Section 2.3) we conducted an unpaired t-test to determine whether there were differences in firing frequency (Hz) and sEPSC amplitude (pA) as a function of PHTPP application. For patch-clamp electrophysiology in rats, we conducted an unpaired t-test to determine whether percent change in DA cell firing frequency following DPN application from a pre-DPN baseline was altered between control and ERβ knockdown rats. Significance was set at *p*<0.05.

#### SA

All SA statistical analyses were conducted using R statistical Program (R Core Team, 2023). Linear mixed effects (LME) were fit in the *lme4* package [64], and where appropriate, post-hoc testing was conducted using *emmeans* (Lenth, 2025). We ran identical LME models on nicotine consumption and discrimination index during nicotine SA. We included random effects of subject and session to account for the interdependence of our repeated measurements that violate analysis of variance (ANOVA) assumptions [65]. Session was included as a continuous factor, and virus group was included as a between-subjects variable. The session by virus interaction was also included in the model to determine whether the slope of the session effect significantly varied by virus group, and whether consumption/discrimination varied between virus groups as sessions progressed. The control virus group was the reference group in all analyses unless otherwise specified for post-hoc testing. Where appropriate, slopes and/or simple effects were probed post-hoc and corrected for multiple comparisons using Tukey’s correction. Significance was set at *p*<0.05. Only the first 11 sessions of SA were included in these analyses due to loss of catheter patency at later sessions. All rats completed a minimum of 11 sessions with patent catheters.

## 3. Results

### 3.1. ERβ Antagonism Reduces VTA DA Excitability in Mice

To determine if ERβ pharmacological antagonism alters DA neurophysiology, we first patched DA neurons within the VTA of ovary-intact female mice and bath applied PHTPP. We conducted an unpaired t-test on firing frequency (hZ) and firing amplitude (pA) between PHTPP (+) and non PHTPP (-) slices (**Figures 1A and 1B**). While there were no effects of PHTPP on firing frequency (p=0.610; **Figure 1E**), rats given PHTPP exhibited significantly lower firing amplitude (p<0.05; **Figure 1F**).

### 3.2 DA ERβ Knockdown Within the VTA Reduces DA Firing Frequency

We show that our AAV (see construct map in **Figure 2A**) successfully infects TH^+^ neurons within the VTA (**Figure 2B**). We also show that GFP^+^ neurons infected with our AAV construct can be readily identified and patched for whole cell electrophysiology (**Figure 2C**). Next, we tested if our ERβ knockdown manipulation successfully impacted DA neurophysiology. We conducted an unpaired t-test on percent change from baseline (before DPN application) for DA cell firing frequency in slices applied with DPN between control and ERβ Knockdown rats. Control rats exhibited significantly greater increases from baseline DA firing frequency relative to ERβ Knockdown rats, t(4)=2.973, p=0.410; **Figures 2D and 2E**.

### 3.3 ERβ Knockdown Within VTA DA Neurons Reduces Consumption During Nicotine SA Acquisition

There were no group differences in food training prior to nicotine SA (**Figure 3B**; *p*>0.05), indicating that ERβ knockdown in VTA DA neurons had no impact on operant food SA. For our LME examining nicotine consumption (**Figure 3C**), we found a significant effect of session β=0.134, t(1,14)=2.354, indicating that control rats increased their nicotine consumption by 0.134 mg/kg per session. We also found a significant Session by Virus interaction, β=-0.199, t(1,14)=-2.471, p=.027, suggesting that the rate of increase in consumption over sessions was attenuated in ERβ knockdown rats by 0.199 mg/kg per session. Thus, the estimated slope of the effect of session on consumption for ERβ knockdown rats was –0.065 mg/kg per session. To determine whether the slope of the effect of session was significant for ERβ knockdown rats (e.g., did consumption significant decrease over time), we re-ran the model with the ERβ knockdown group as the reference group. Notably, consumption did not significantly change over time for the ERβ knockdown rats, β=–0.065 mg/kg, t(1,14)=-1.141, p=0.273. Additional post-hoc testing examining pairwise differences in consumption between control and ERβ knockdown rats at each session determined that consumption was estimated to be significantly greater in control rats relative to ERβ knockdown rats for sessions 9-11 (ps≤.029). In sum, consumption increased over time in the control virus group but did not change over time in the ERβ knockdown group, such that by the end of SA, the ERβ knockdown group consumed less nicotine than the control virus group (**Figure 3C**).

### 3.4. ERβ Knockdown Within VTA DA Neurons Does Not Impair Lever Discrimination During Nicotine SA

We next performed this same LME model on lever discrimination (Figure 3D). There were no significant effects of Session [β=-0.005, t(14)=-0.686, p=0.504], Virus Group [β=-0.067, t(14)=-0.766, p=0.456] nor was there a Session by Virus Group interaction [β=0.009, t(14)=0.735, p=0.475]. Thus, even though nicotine consumption was suppressed in the ERβ knockdown group relative to the control virus group (see Section 3.3), rats did not differ in their lever discrimination (**Figure 3D**). These results indicate that ERβ knockdown within VTA DA neurons *specifically* decreased nicotine consumption without impairing learning of the lever discrimination, similar to our prior results following OVX[27; 43].

## Discussion

Here we show that akin to OVX[38; 43], ERβ knockdown in VTA DA neurons results in significant decreases in nicotine consumption during acquisition of SA in ovary-intact female rats. These results are important because they are the first to demonstrate reductions in nicotine use due to a cell-type specific mechanistic interaction between estrogenic signaling and the mesolimbic DA system, which can recapitulate effects on nicotine SA found when endogenous sex hormone production and release is abruptly halted via OVX [66]. We further show that the reduction in nicotine consumption by ERβ knockdown in VTA DA neurons was *specific* to nicotine, as no differences were found in food training prior to nicotine SA acquisition. As well, differences in nicotine SA acquisition cannot be attributed to baseline differences in operant lever press acquisition given the lack of group differences in food training. Here we also developed a novel AAV construct which allows for knockdown of ERβ in VTA DA neurons, which has functional impacts on DA signaling. Importantly, these results demonstrate an outsized role of estrogenic signaling on DA neurophysiology and nicotine use. We have previously shown that OVX downregulates VTA ERβ protein expression in nicotine self-administering females, however, our prior study did not demonstrate that this downregulation caused the decreased nicotine consumption observed in OVX females [43]. While other mechanisms may still be involved, our current results suggest that OVX-induced VTA ERβ knockdown may have been an important mechanism underlying the decreased nicotine use associated with OVX in females.

While our results fill a gap in our understanding of female-specific mechanisms driving nicotine use acquisition, it remains unclear if ERβ plays a similar role in gating nicotine SA acquisition in males. Indeed, our results may underscore a potential latent sex difference in how ERs regulate DAergic mechanisms underlying nicotine use, although studies supporting this possibility need to be done. Indeed, there are no studies to date that have evaluated ER-related mechanisms driving nicotine use in males. Somewhat surprisingly given that wealth of literature on the neuroscience of nicotine addiction that exclusively evaluated male subjects across decades, there is less known regarding how gonad-derived steroid hormones impact male nicotine use as this is rarely evaluated in preclinical nicotine SA models. Only one study to our knowledge has evaluated hormone contributions to nicotine SA in adolescent male rats and found no effects [39] but gonadectomy was not conducted and thus it is difficult to determine if the outcomes in this study were due only to testosterone and not to other steroid hormones including E2. Despite this lack of rigor of prior research, there is some empirical support that steroid hormones sex-specifically impact female addiction neurobiology. One early study found that while OVX and E2 replacement significantly modulated extracellular striatal DA in females, this manipulation did not alter dopamine levels in males [44] indicating sexually dimorphic impacts of gonad-derived hormone loss on mesolimbic DA signaling. Bringing these effects into focus with addictive drugs, one study found that basal and methamphetamine-induced extracellular DA in the striatum was reduced in OVX female rats, which was restored by E2 supplementation [67]. Thus, while a significant gap in our understanding of sex differences in DA regulation of nicotine use remains, there are prior studies indicating that there may be sex-specificity within the mesolimbic dopamine circuit which is heavily implicated in nicotine SA [29; 68; 69].

As noted above, we have created a novel AAV that we confirm functionally reduces ERβ regulation of VTA DA neurons. We have further confirmed that the viral construct infects TH^+^ neurons within the VTA. We developed this construct because while lentiviral constructs have been developed that allow for knockdown of ERα or ERβ in the VTA [70], these constructs do so without cell-type specificity limiting interpretation of how ERs may regulate DA signaling. Further, this prior construct only works in mice, thus limiting its use in rat SA paradigms. Given our current data in Figure 1, prior studies indicating that E2 regulates VTA DA neuronal excitability, and that the VTA is a heterogenous brain region [26; 47], determining how ERs specifically located on DA neurons within the VTA is critical to our understanding of female-specific nicotine neurobiology.

Limitations to our study include a focus on females without similar evaluations in males. Further, here we only tested nicotine SA acquisition at one unit dose, and thus future studies are needed to determine if VTA ERβ knockdown-induced suppression of nicotine consumption during acquisition is dose-dependent. An additional limitation to our studies is that we did not evaluate the impacts of ERα knockdown in VTA DA neurons on nicotine use acquisition, which may also play an important role in driving nicotine use given our prior study demonstrating that nicotine SA suppresses the ability of E2 to increase transcription of VTA ERα following OVX in female rats. We have developed a similar AAV to specifically knockdown ERα in VTA DA neurons and our future studies will validate this knockdown to evaluate whether ERα on VTA DA receptors play a commensurate role in altering suppression. Future studies should also evaluate the role of progesterone in DAergic signaling underlying nicotine use, given clinical studies showing that progesterone reduces nicotine use in women [71; 72; 73]. Finally, while it is unlikely that our specific knockdown of ERβ on VTA DA cells altered cyclicity, here we did not track estrous cycle phase. Thus, future studies are needed to determine if cell-specific knockdown of ERβ within the reward pathway has larger effects on the reproductive system. Notably, this is an unlikely outcome given that E2 regulates the estrous cycle via ERs in hypothalamus and influences signaling between the hypothalamus, the pituitary gland, and the ovaries (the hypothalamic-pituitary-gonadal axis; [74]. Thus, unsurprisingly, there is no established role of VTA DA neurons in regulation of hormonal cycling and we would not expect our cell-specific ERβ knockdown manipulation to have impacted cyclicity.

## Conclusions and Future Directions

Here we show that VTA DA neurons are critically regulated by ERβ, and this drives nicotine use acquisition in ovary-intact female rats. Our results have important implications for our understanding of female-specific nicotine neurobiology, as clinical reports indicate that ovary-derived steroid hormones play an important role in women’s smoking patterns [17; 18; 19]. Our results extend prior studies utilizing OVX by evaluating cell-type specific mechanisms of nicotine neurobehavior in ovary-intact female rats. As well, our results further underscore the importance of evaluating nicotine neurobehavior through the lens of sex as a biological variable, as there may be latent sex differences in smoking trajectories that are not well understood. Future studies are needed to evaluate how ERβ-induced regulation of nicotine use acquisition may differ as a function of reproductive aging, evaluating these female-specific mechanisms across critical reproductive transitions including sexual maturity during puberty and menopause. Indeed, little is known regarding female nicotine neurobiology and behavior across critical reproductive aging windows, highlighting an important gap in the rigor of prior research and underscoring the need for future studies to evaluate these relationships.

**Table 1.** Antibodies and dilutions for immunohistochemistry.

| Antibody Name | Immunogen | Antibody Info | Dilution |
| --- | --- | --- | --- |
| Anti-tyrosine hydroxylase (TH) | Two synthetic peptide/keyhole limpet hemocyanin (KLH) conjugates | Abcam; catalog #AB76442; RRID:AB_1524535 chicken (polyclonal) | 1:2000 |
| Alexa Flour 594 anti-rabbit IgG |  | ThermoFisher Scientific; catalog #A-11012 | 1:500 |
| Alexa Flour 488 anti-chicken IgG |  | ThermoFisher Scientific; catalog #A-11039 | 1:500 |

